# Heterogeneous Sensitivity to Src Inhibitors in Oral Squamous Cell Carcinoma and Its Implications for Combination Therapy with Cisplatin

**DOI:** 10.64898/2026.04.02.716058

**Authors:** Yuri Ofusa, Tadahide Noguchi, Hiroaki Mizukami, Kenji Ohba

## Abstract

Treatment options for advanced oral squamous cell carcinoma (OSCC) are limited, and cisplatin toxicity and drug resistance are major clinical challenges. Src is a central kinase that integrates multiple oncogenic pathways and a promising therapeutic target; however, Src inhibitors have shown limited efficacy as monotherapies and their sensitivity in OSCC remains elusive. Herein, we examined the activation of major oncogenic signaling pathways and the antitumor effects of FDA-approved Src inhibitors *in vitro* and *in vivo* using multiple OSCC cell lines and a mouse model. The activation of the key pathways, including Src and MAPK, considerably differed among the cell lines and was linked to heterogeneous sensitivity to Src inhibitors. Effective growth suppression required Src dephosphorylation and downstream MAPK pathway inhibition, which vary depending on the cell line. Additionally, combination treatment with an Src inhibitor and cisplatin showed additive antitumor effects, facilitating the decrease of cisplatin doses by half without efficacy loss. Notably, dasatinib alone and in combination with cisplatin decreased tumor burden with characteristic internal tumor death *in vivo*. These results elucidate OSCC dependency on Src signaling and the potential of Src inhibition to decrease cisplatin toxicity, thereby paving way for new therapeutic strategies.

## Introduction

Oral squamous cell carcinoma (OSCC) is a malignant disease on the mucosal surfaces of the tongue, gingiva, and lips, and is the most common subtype of head and neck malignancy^1^. Approximately 390,000 new cases of OSCC were reported worldwide in 2022, and the rate of increase is predicted to reach nearly 60% by 2050, according to the Global Cancer Observatory ^2^.

Almost 60% of patients with oral cancer are diagnosed at advanced stages (stage III–IV), with a 5-year survival rate of less 60%^3–6^. Managing patients with advanced cancer requires multimodal treatment including surgery, radiotherapy, and chemotherapy^6^. Although multidisciplinary therapy is effective for managing advanced cancer, it is quite invasive, resulting in treatment-related cosmetic and functional impairments such as dysphagia and speech difficulties, which considerably compromise the quality of life(QOL) patients^7^. Moreover, 10–50% of patients experience local recurrence or distant metastasis following initial therapy, and only a minority are eligible for salvage surgery or radiotherapy^8–10^. Palliative or supportive care is the last option for most patients with recurrent or metastatic cancer, with a median survival of 4–9 months^3,11^. Thus, improving the prognosis and QOL of patients with OSCC remains an urgent clinical challenge.

Systemic chemotherapy plays a key role, especially in patients with recurrent or metastatic cancer. Cisplatin is the standard systemic therapy in definitive chemoradiotherapy and recurrent/metastatic settings. However, owing to severe toxicities, few patients with adequate performance status (PS) can be prescribed cisplatin^12^. No alternative systemic therapy with efficacy comparable to that of cisplatin has been established for patients ineligible due to age, comorbidities, or poor PS^3^. In addition, cisplatin resistance occurs in approximately 30–40% of patients with OSCC, substantially limiting its therapeutic effect^13^. Therefore, drugs that decrease cytotoxicity and overcome cisplatin resistance are necessary.

Several studies have explored the use of cisplatin substitutes. While carboplatin shows lower toxicity than cisplatin, its antitumor activity is inferior to cisplatin at equivalent doses^14^. Cetuximab is the only molecular-targeted drug approved for OSCC. However, it has not shown clear superiority over cisplatin yet^15^. Even though immune checkpoint inhibitors such as pembrolizumab and nivolumab have recently been approved for treating recurrent/metastatic OSCC, their response rates are limited to approximately 10–20%^16,17^. The identification of new effective therapeutic agents for OSCC is a crucial unmet clinical need.

A central non-receptor tyrosine kinase within the Src family of kinases (SFKs), c-Src (hereafter, Src), regulates multiple cellular processes involved in malignant progression in tumors, including proliferation, survival, migration, invasion and angiogenesis^18^. Src hyperactivation has been noted in solid tumors, including breast, colorectal, pancreatic, ovarian, gastric, esophageal, and lung cancers and is strongly liked to poor prognosis^19^. This indicates that Src inhibitors are potential candidates for cancer chemotherapy. Src activation is correlated with poor clinical outcomes and invasive and metastatic potential in head and neck squamous cell carcinoma (HNSCC)^20^. Particularly, in OSCC, high Src expression is markedly linked to tumor size, lymph node metastasis, advanced stage and recurrence, and has been identified as an independent prognostic factor. This suggests that Src plays a crucial role in OSCC progression^21^.

Preclinical studies have revealed that FDA-approved Src inhibitors, including dasatinib and saracatinib, suppress proliferation, migration and invasion, and induce apoptosis in HNSCC/OSCC models^19,22,23^. Despite these promising results, Src inhibitors have demonstrated suboptimal efficacy as monotherapies in clinical trials^24,25^. Possible mechanisms behind this discrepancy include the non-lethal nature of Src inhibition in HNSCC/OSCC cells, compensatory activation of alternative pathways, such as PI3K/AKT and STAT3^22,26^, and variations in EMT status or cell adhesion molecules that affect drug sensitivity^27,28^. Substantial efforts have been directed toward elucidating the mechanisms of action of Src inhibitors and identifying biomarkers that predict the therapeutic response in lung, colorectal, and breast cancers, where oncogenic driver mutations are well characterized. However, the molecular determinants of Src inhibitor sensitivity in OSCC remain elusive. HNSCC/OSCC is characterized by profound molecular heterogeneity, with distinct dependencies on kinases, such as EGFR, EPHA2, AURKA, WEE1, and JAK1 across various cell lines^29^. This heterogeneity suggests that sensitivity to Src inhibition may markedly differ among the OSCCs. However, there is no report systematically investigating Src inhibitor sensitivity in OSCC, especially using animal models. Moreover, recent the combined effects of Src inhibitors and cisplatin may provide a strategy to overcome cisplatin resistance and decrease toxicity in gastric cancer and tubular epithelial cells^30,31^.

Based on this evidence, we sought to reassess the therapeutic potential of Src inhibition in OSCC and generate foundational insights that may guide the optimization of future treatment strategies. We analyzed Src inhibitor sensitivity across multiple OSCC cell lines and a mouse model and assessed the therapeutic effect of combining Src inhibition with cisplatin.

## Results

### OSCC cell lines show substantial heterogeneity in the activation of major oncogenic signaling pathways

Cancer develops through the accumulation of genetic and epigenetic changes in oncogenic signaling pathways, eventually resulting in the acquisition of malignant phenotypes^32^. In OSCC, mutations frequently occur in p53 gene^33^, and multiple oncogenic pathways including EGFR, PI3K/AKT/mTOR, JAK/STAT, MET, Wnt/β-catenin and RAS/RAF/MAPK are aberrantly activated and contribute to tumor progression^34^. Besides, the activation of MAPK signaling (ERK1/2, JNK, p38, and ERK5) is involved in HNSCC oncogenicity^34^. These pieces of evidence suggest that the abnormal activation of signaling networks should be required, but the activating status may markedly vary in the OSCC cell lines.

To test this hypothesis, we investigated the phosphorylation and activation status of key signaling molecules for p53, Src, Ras and MAPK by western blotting (Fig. 1A). The increased proteins and phosphorylation levels of p53 (p-p53; Ser15) were observed in HSC-4 and CAL-27 cells. Total Src expression was consistent in all cell lines except SCC-25. Otherwise, the phospho-Src (p-Src; Tyr416), which is active form, was detected in all cell lines although the activated level differs (high: SAS Ho-1-u-1 and CAL-27, modest: HSC-2 and HSC-4, low: HSC-3 and SCC-25). K-Ras expression was augmented in SAS, HO-1-u-1, and CAL-27, and the discrepancy between K-Ras and total Ras expression in HSC-2 and SAS suggested N-Ras or H-Ras activation. In the MAPK pathways, the increase of total p38, Erk1/2 and SAPK/JNK expression seemed to be corresponding to the activation level of Src in all cell lines except HSC-3. The strong phosphorylation of SAPK/JNK (p-SAPK/JNK) was observed in SAS and HO-1-u-1 cells, which showed stronger activation of Src compared to other cell lines. Notably, the phosphorylated level of p38 and ERK1/2 (p-p38 and p-ERK1/2) are almost related to activation of Src (p-Src) in all cell lines. This concordant activation pattern suggested that Src activity was closely associated with downstream MAPK signaling, especially about p38 and Erk1/2, in the most of OSCC cells. Thus, as different from the case of p53 and Ras randomly activated (the case of random activation in p53 and Ras), broadly activated Src followed by the activation of MAPK signals may be a common oncogenic pathway in most OSCC cell lines, meaning that (raising the possibility that) Src could be a promising target for chemotherapy in OSCCs.

**Figure 1.**
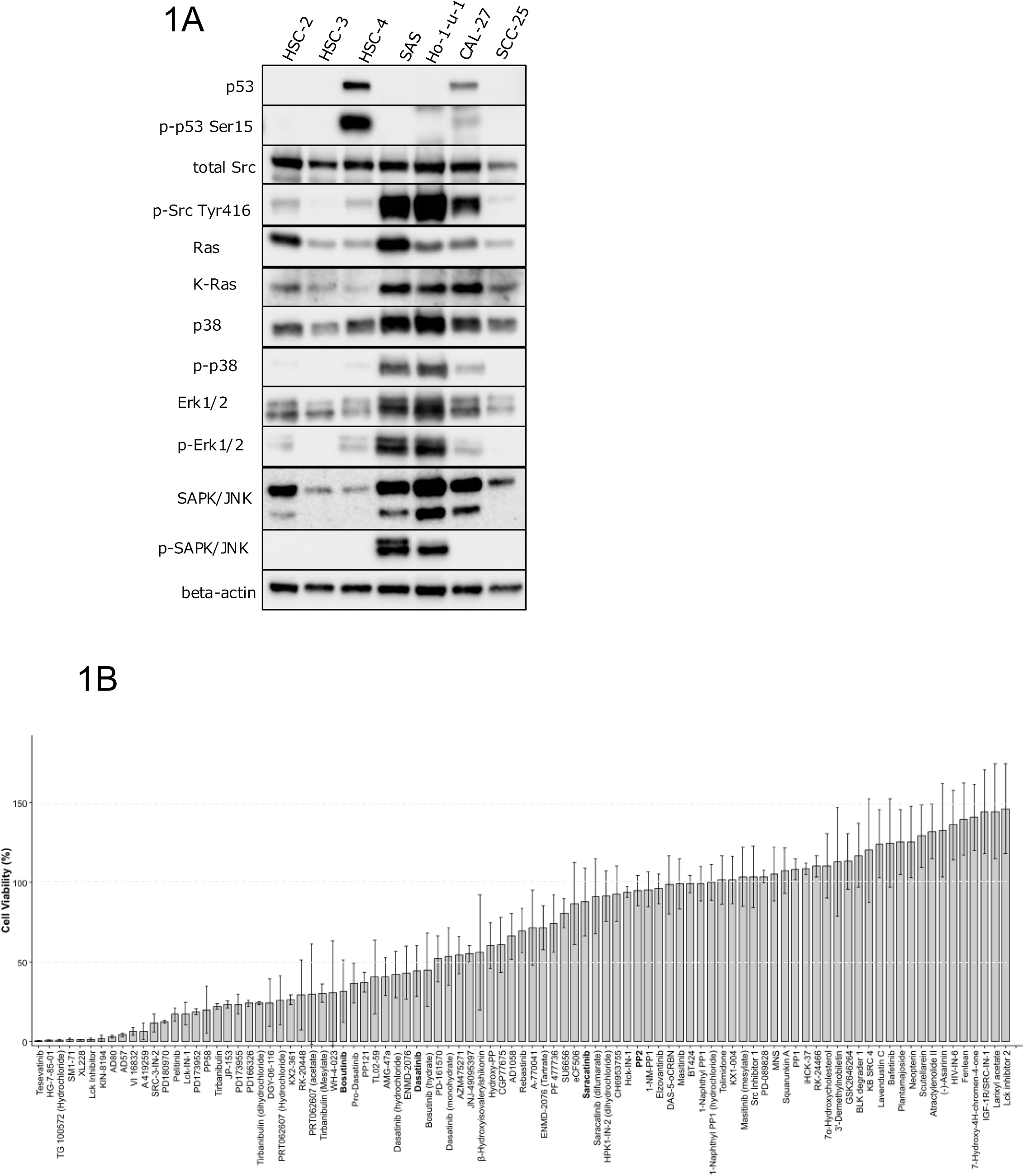
Activation of oncogenic pathways in OSCC cells and screening of a 93 Src-associated chemical compounds. **A** Western blot analysis of the oral squamous cell carcinoma (OSCC) cell lines to monitor numerous signaling pathways. The phosphorylated and total protein levels were determined as indicated. **B** Screening with a library of 93 Src-associated chemical compounds(10µM) was performed using CAL-27 cells. Cells were treated for 36 h, and viability was measured using the CellTiter-Glo assay and normalized to the 1% dimethyl sulfoxide (DMSO) control. Data represent mean ± standard deviation (SD) from three independent experiments (n = 3).

### Src activity and its inhibitory effect of tumor growth by specific inhibitors in different OSCC cell lines

Our data as shown in Fig. 1A and the fact that some Src inhibitors have already been approved for clinical cancer therapies^19,22,23^ suggest that these Src inhibitors can be easily utilized for OSCC if their anticancer effect is prospective. To narrow down promising candidates, we initially screened 93 Src-associated compounds in CAL-27 (Fig. 1B, Table S3) which show the moderate activation of all oncogenic pathways such as p53, Ras and Src. The data in Fig. 1B indicates that at least half of Src inhibitors showed tumor growth suppression (Viability <100 %). Among these compounds, we focused on FDA-approved Src inhibitors, which have not been clinically applied in HNSCC/OSCC yet, allowing for more rapid clinical translation because of well-established safety profiles and lowers regulatory barriers. Next, we evaluated the sensitivity of the OSCC cell lines to FDA-approved Src inhibitors. Despite PP2 is not FDA-approved and ponatinib and sarcatinib are not listed in initial screening, these were included for comparison because it is frequently used as a research-grade and FDA-approved Src inhibitor, respectively. Treatment of seven OSCC cell lines with each Src inhibitor and cisplatin which is prestigious tumor growth inhibitor as comparison showed variation in cell line viability (Fig. 2A). HSC-2 and SCC-25 were broadly sensitive to all the inhibitors tested, whereas SAS showed weak sensitivity to various Src inhibitors except bosutinib (Fig. 2A) although the highest susceptibility was observed to cisplatin. Comparison of the IC₅₀ values obtained following 36 h of treatment indicated differences between the inhibitors (Table 1). Dasatinib, vandetanib and saracatinib revealed pronounced variability in their IC50 values across cell lines. In contrast, ponatinib and bosutinib showed broad antitumor effects in various cell lines (Table 1). Notably, HO-1-u-1 showed resistance to cisplatin. Thus, OSCC cell lines show distinct pharmacological sensitivities resulting from the differences in the activating pathways for Src signaling.

**Figure 2.**
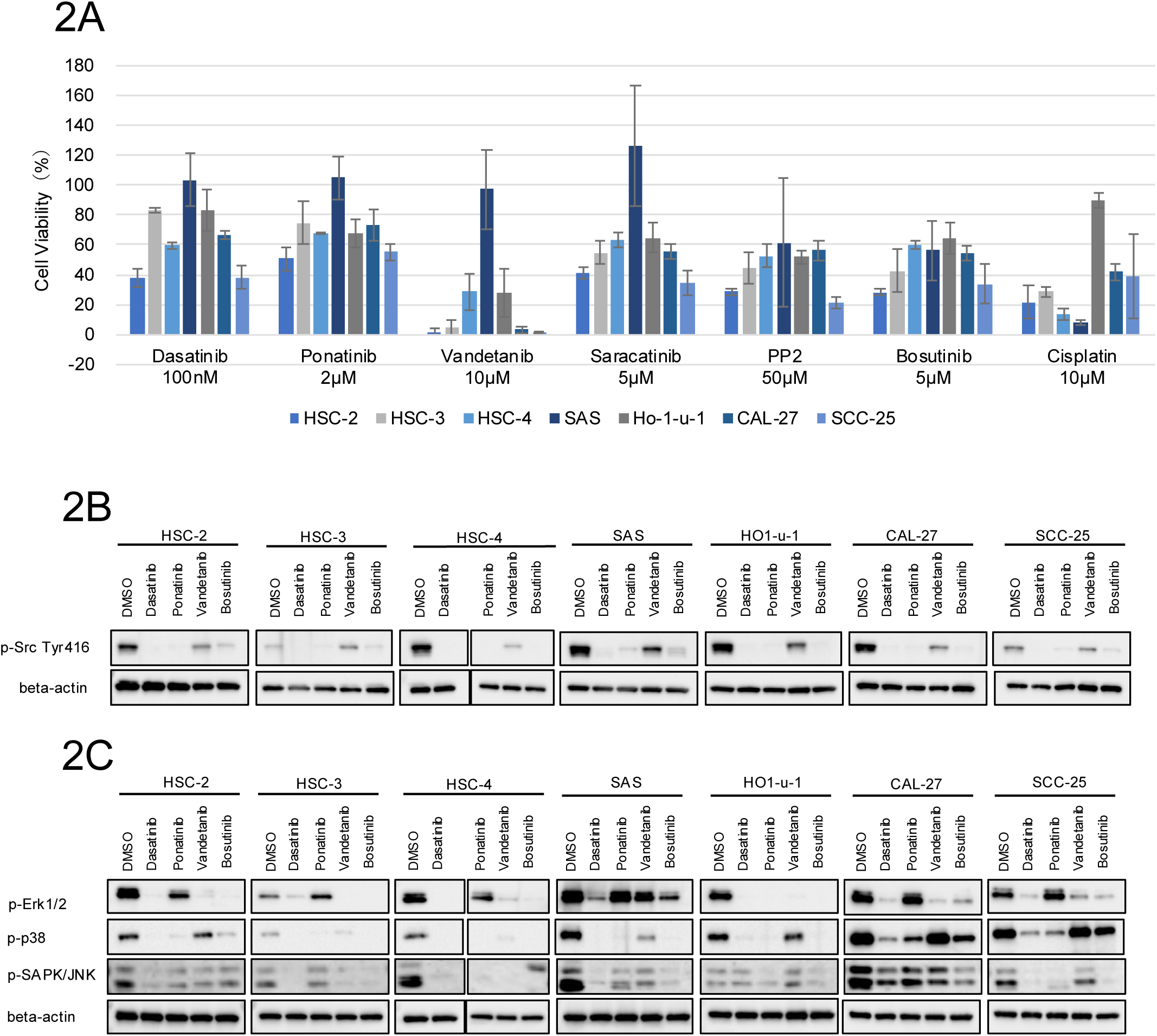
Antitumor effects of FDA-approved Src inhibitors in various OSCC cell lines and their impact on Src activity. **A** Cell viability (%) of the OSCC cell lines treated with each Src inhibitor for 72 h. Viability was measured using CellTiter-Glo, and values are indicated relative to the 1% DMSO control. Data represent mean ± SD from three independent experiments (n = 3). **B**,**C** Western blot analysis of the OSCC cell lines to investigate Src (B) and MAPK (C) activity induced by different Src inhibitors. OSCC cell lines were treated with an Src inhibitor for 18 h, followed by western blot analysis. B indicates the Src phosphorylation (Tyr416), and C reveals MAPK pathway-related proteins. Supplemental Table S4 lists the concentrations of the Src inhibitors used.

**Table 1.** The half maximal inhibitory concentration (IC50) of Src inhibitors in various OSCC cell lines after 36 h of treatment.

|  | Dasatinib | Ponatinib | Vandetanib | Saracatinib | PP2 | Bosutinib | Cisplatin |
| --- | --- | --- | --- | --- | --- | --- | --- |
| HSC-2 | 0.1 μM | 2 μM | 5 μM | 5 μM | 50 μM | 5 μM | 10 μM |
| HSC-3 | 10 μM | 2 μM | 8 μM | 10 μM | 100~250 μM | 5 μM | 1~5 μM |
| HSC-4 | 25 μM | 5~10 μM | 25 μM | 100 μM | 600 μM | 15 μM | 10 μM |
| SAS | 10 μM | 5~10 μM | 50 μM | 100 μM | > 600 μM | 25 μM | 10 μM |
| Ho-1-u-1 | 50 μM | 5~10 μM | 25 μM | 50~100 μM | > 600 μM | 25 μM | > 50 μM |
| CAL27 | 10~25 μM | 5~10 μM | 10~50 μM | 25 μM | > 600 μM | 15 μM | 10 μM |
| SCC-25 | 10 μM | 2~5 μM | 8~10 μM | 50 μM | 200 μM | 10 μM | 10 μM |

We next examined the level of activated Src (p-SrcTyr416) levels following inhibitor treatment to investigate the relativity between the antitumor activity and the degree of Src inhibition (Fig. 2B). All inhibitors suppressed Src activation, although vandetanib relatively exhibited a weaker inhibition in most of cells and no inhibition in HSC-3. On the other hand, SAS showed weaker suppression of Src activation by ponatinib, vandetanib, and bosutinib compared with the other cell lines, which is consistent with their reduced sensitivity on growth suppression (Figs. 2A).

We further assessed whether Src inhibition was correlated with the suppression of the MAPK subpathways (ERK1/2, p38, and JNK; Fig. 2C) in OSCC cell lines. Dasatinib strongly reduced the most of activated MAPK components in all the cell lines except SAPK/JNK in HO-1-u-1. Bosutinib typically inhibited the phosphorylation of all MAPK components, although the inhibitory effect for ERK1/2 in SAS, and p38 in CAL-27 and SCC-25 were weak or not observed, respectively. Ponatinib robustly and broadly inhibited the activation of p38 in all cel lines, whereas JNK inhibition was slightly limited in CAL-27 and not observed in HSC-3. The relative inhibitory level of Erk1/2 by ponatinib was no or weak in various cells such HSC-3, SAS, CAL-27 and SCC-25. Vandetanib showed the random pattern of MAPK pathways inhibition in cell lines used. Thus, while the downstream MAPK pathways activated Src varies in cells and the strength of inhibitory effect is different, inhibitors targeting Src activity, especially dasatinib and bosutinib, are likely effective to suppress the growth of OSCC cell lines.

Taken together, these results suggest that although the MAPK signaling pathways related to tumorigenesis significantly differs across the OSCCs, the Src, which has the central role for MAPK activation at upstream signaling, could be a feasible candidate as anti-tumor drugs.

### Tumor inhibitory effect by combination therapy using cisplatin and Src inhibitors

OSCC cell lines showed heterogeneity such as variable susceptibilities to Src inhibitors (Fig. 2). Next, we assessed the combinatorial effects of Src inhibitors with cisplatin (Fig. 3). All Src inhibitors have higher suppressive effect than cisplatin at 5µM on tumor growth. Compared with cisplatin at a clinically relevant dose of 10 µM, Src inhibitors alone showed comparable or slightly lower antitumor activity in HSC-2, HSC-4, SAS, and CAL-27. HSC-3 treated with Src inhibitors exhibited lower antitumor effects than those treated with cisplatin (10µM), while SCC-25 and HO-1-u-1 showed higher antitumor activity in response to Src inhibitors than to cisplatin (10µM). This indicates that Src inhibition was almost as effective as cisplatin, the standard treatment for OSCC. The combination treatment of cisplatin (5µM) and each Src inhibitor showed additive effects, which indicate that the combined antitumor activity is approximately equal to the sum of the effects observed with each agent alone, in HSC-3, HSC-4, CAL-27, and SCC-25. This effect was especially pronounced with dasatinib and bosutinib because the observed cell viability levels were close to the combined ratios calculated from the single inhibitor treatments. Although cisplatin at 5 µM alone showed limited activity, its combination with Src inhibitors produced antitumor effects comparable to or more for 10 µM cisplatin alone in HSC-4, CAL-27, and SCC-25. On the other hand, cisplatin-resistant HO-1-u-1 responded better to Src inhibitors alone than to cisplatin, and no enhancement was noted with the combination treatment. This indicates that the efficacy of Src-cisplatin combination therapy may be determined by the intrinsic cisplatin sensitivity of each OSCC cell line and its degree of dependence on Src signaling. These results suggest that combining Src inhibitors with cisplatin may reduce the cisplatin dose to half of the standard *in vitro* therapeutic concentration while maintaining its efficacy in cisplatin-sensitive OSCC.

**Figure 3.**
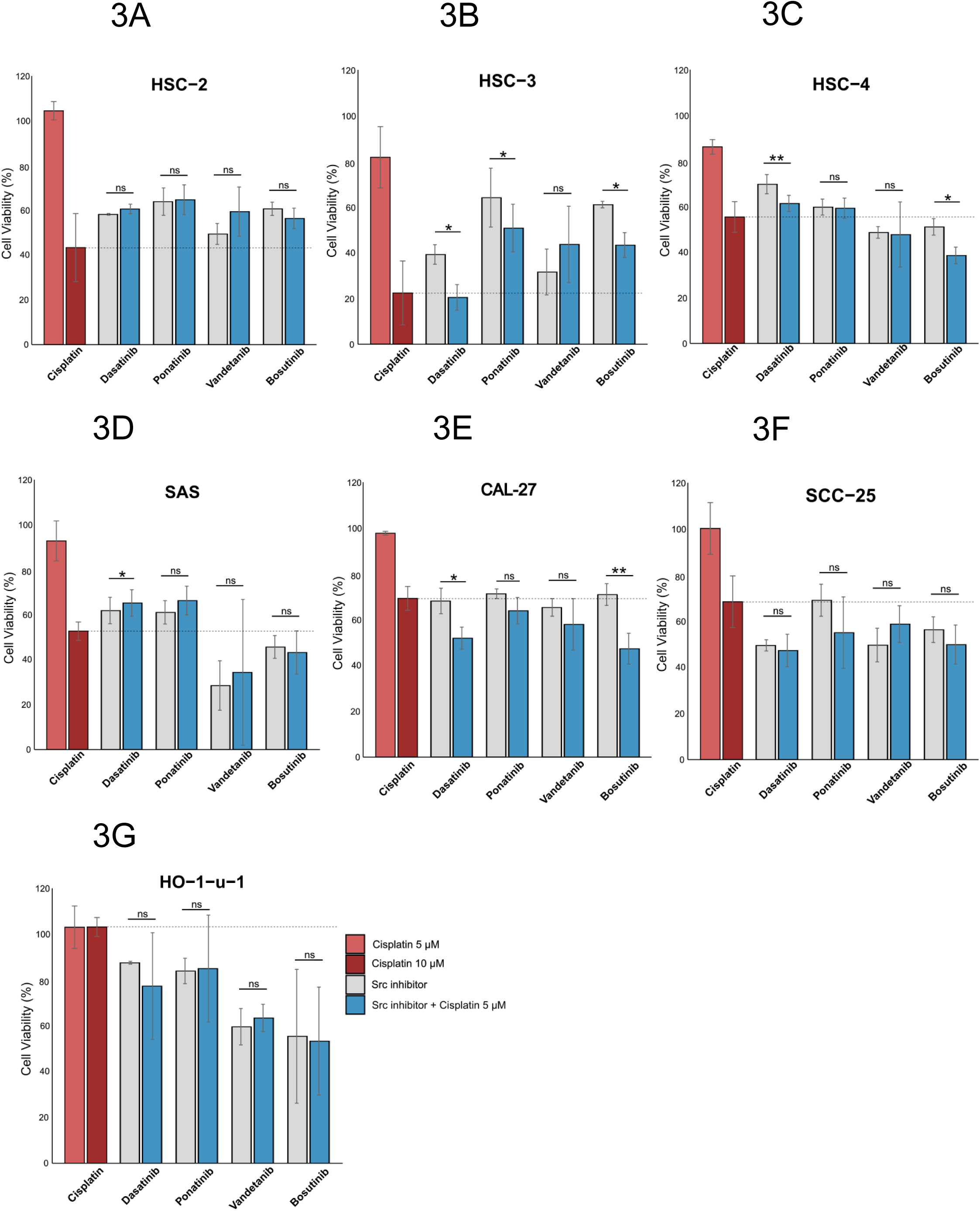
The tumor inhibitory effect by combination therapy using cisplatin and Src inhibitor. Cells were treated with cisplatin (10 µM or 5 µM), each Src inhibitor (dasatinib, ponatinib, vandetanib, or bosutinib) alone, and the combination of cisplatin (5µM) with each Src inhibitor for 36 h. Cell viability is indicated as the percentage relative to the 1% DMSO control. Data represent mean ± SD from three independent experiments. A–G show the data for the indicated cells. P-values were calculated using paired t-tests in Microsoft Excel. Statistical significance was indicated as p < 0.01 **, p < 0.05 *, and p ≧ 0.05 ns.

### Tumor growth inhibition by single treatment of dasatinib and bosutinib *in vivo*

Dasatinib and bosutinib showed strong inhibitory effect on Src activation with following consistent suppression of MAPK signaling pathways in most of OSCC cell lines compared to other Src inhibitors. In addition, CAL-27 showed that moderate level of activating oncogenic signals (Src, p53 and Ras) among cell lines. Since dasatinib and bosutinib demonstrated additive effects with cisplatin in more cell lines than the other Src inhibitors (Fig. 3) and both are relatively inexpensive, thus more suitable for clinical applications. Consequently, we determined to use dasatinib and bosutinib as Src inhibitors, and CAL-27, which showed reliable tumor formation in preliminary experiments, as model of OSCC for further *in vivo* analysis.

We first examined whether dasatinib and bosutinib, which showed additive effects *in vitro*, could suppress tumor growth of CAL-27 as single agent *in vivo*. To monitor tumor size using *in vivo* imaging system, we established stable Rluc-expressing CAL-27 (CAL27-Rluc) cells and checked their susceptibility to Src inhibitors (Fig. S1). Src inhibitors suppressed the growth of CAL27-Rluc clones in a similar manner as the CAL-27 Wt cells. Clone7, which exhibited a median susceptibility to Src inhibitors, was selected for *in vivo* analysis.

CAL27-Rluc cells-inoculated BALB/cAJcl nu/nu mice were treated with dasatinib (30 mg/kg, Fig. 4 and Fig. S2) or bosutinib (50 mg/kg, Fig. S3) by the indicated schedule (Fig.4A and Fig. S3A) which were designed to approximate clinically relevant regimens. Cisplatin 4 mg/kg was selected as a positive control based on previous reports, and this dose is within the range of the maximum repeated tolerated dose in the mouse model^35^.

**Figure 4.**
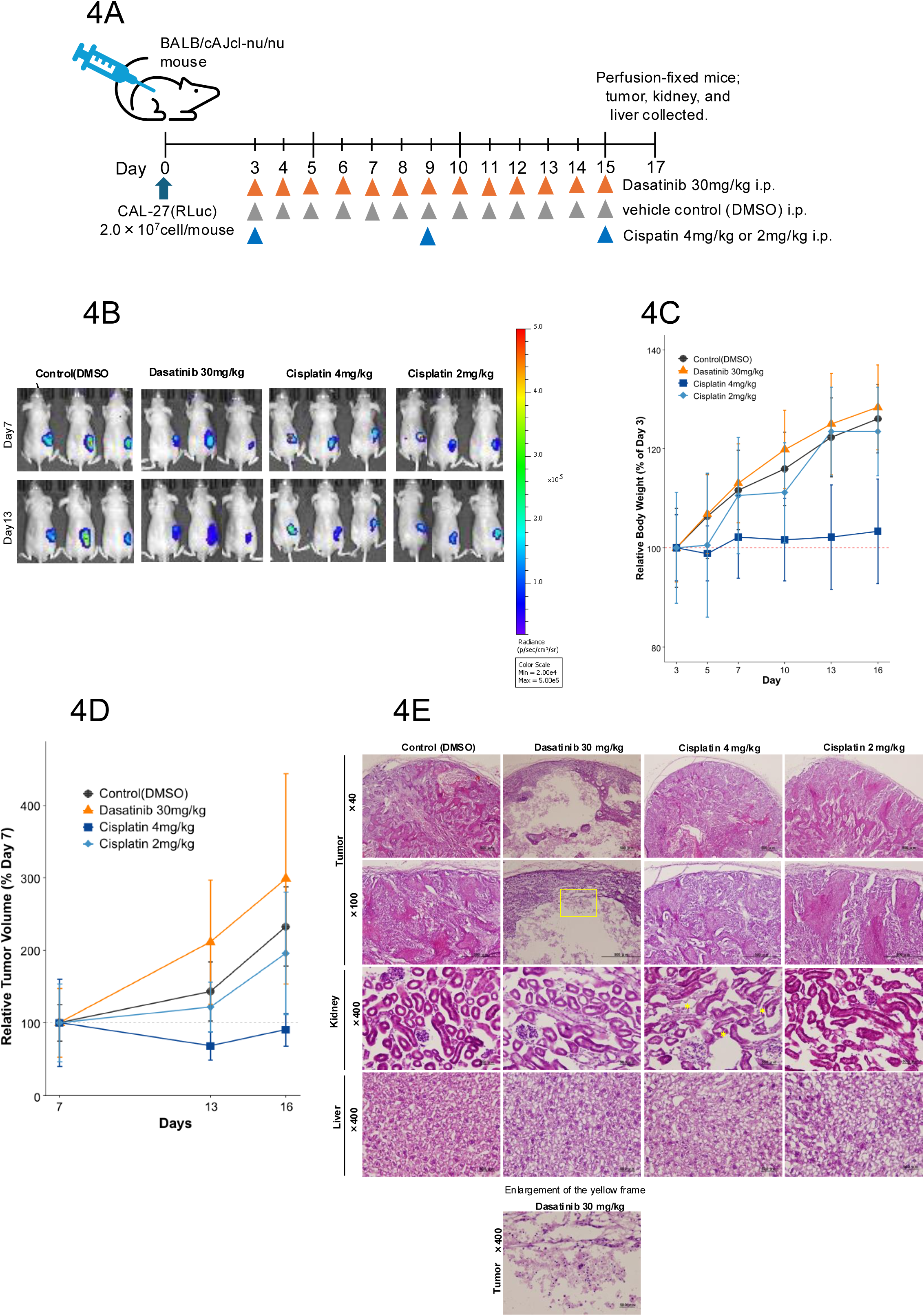
Tumor growth inhibition by single treatment of dasatinib in vivo. **A** Experimental schedule for a single treatment with dasatinib *in vivo*. RLuc-expressing CAL-27 (CAL27-Rluc) cells (clone 7) (2 × 10^7^ cells/mouse) were subcutaneously implanted into the right flank of 5-week-old male BALB/cAJcl nu/nu mice. Three days following implantation, the mice were randomly assigned to four treatment groups: vehicle control (DMSO), dasatinib alone (30 mg/kg), cisplatin alone (4 mg/kg), and cisplatin alone (2 mg/kg) (n=5–6). **B** IVIS bioluminescence live imaging of the tumor burden (additional mice are indicated in Fig. S2). Bioluminescence signals were quantified 10 min following ViviRen injection according to the manufacturer’s instructions. **C** Body weights of nude mice from various treatment regimens implanted with CAL27-Rluc xenografts. Body weights are presented as mean ± SD (n = 5–6) and were normalized to the weight on Day 3 for each mouse. **D** Tumor volumes in indicated time points (mean ± s.d., n = 5–6). Tumor volumes were measured using calipers and calculated as L × W² × 0.52. Measurements were conducted at day7, 13 and 16. The tumor volumes were normalized to the volume on day 7 for each mouse. **E** Hematoxylin and eosin staining of the tumor, kidney, and liver sections. Mice were perfusion-fixed on day 17 and the harvested tissues were processed to generate frozen sections for histological analysis. Representative images of each organ are presented. Tumor histology: Control (DMSO) and cisplatin groups showed solid squamous cell carcinoma. Dasatinib group exhibited a hollow, cyst-like structure with evidence of cell death within the lumen. Kidney histology: Control (DMSO) and dasatinib groups displayed normal renal architecture. Cisplatin 4 mg/kg group caused tubular epithelial degeneration and atrophy with focal rupture (*), while cisplatin 2 mg/kg groups induced mild tubular epithelial atrophy. Liver histology: All groups showed normal hepatic morphology without detectable abnormalities. Scale bars: 500 µm (×40, ×100); 50 µm (×400).

The IVIS bioluminescence live imaging showed a substantial reduction in the tumor signal in the dasatinib group relative to the control group (Fig. 4B S2 and S3). Intriguingly, it exceeded the effect of cisplatin in some cases (Fig. 4B, Fig. S2) although bosutinib treatment led to antitumor activity; one of the three mice died, and the remaining animals showed weight loss due to considerable toxicity (Fig. S3B–D), indicating that bosutinib may have limiting utility clinically. Body weight gain was strikingly impaired in the cisplatin 4 mg/kg group, whereas mice treated with dasatinib or cisplatin (2 mg/kg) maintained stable weight without apparent toxicity (Fig. 4C). Curiously, there was a discrepancy between the physical tumor volumes and IVIS imaging data (Fig 4B and D). The physical tumor volumes in the dasatinib group continued to increase compared with the cisplatin 4 mg/kg group (Fig 4D), whereas the IVIS imaging data (Fig. 4B) showed smaller tumor formation in mice treated with dasatinib.

To investigate this discrepancy, histological analyses are performed on day 17. The data showed that tumors from the control and cisplatin groups were solid, well-differentiated squamous cell carcinomas with cancer pearls (Fig. 4E). Contrastingly, tumors from the dasatinib group showed cyst-like structures with hollow interiors and serous fluids. Under magnification, the cells remaining in the lumen had blurred outlines, lost nuclei, and conceivably died. In addition, the cancer pearls in the epithelial regions disappeared. These results indicate that discrepant data between IVIS imaging and physiological tumor size are due to the difference of measuring methods whether the data corresponds contents inside of tumors. In the kidneys, normal kidney structure was maintained in the control group (DMSO) and the dasatinib and bosutinib group (Fig. 4D and S3). On the other hand, degeneration and atrophy of the renal tubular epithelium and localized rupture were observed in the cisplatin 4 mg/kg group, while only mild renal tubular epithelial atrophy was observed in the cisplatin 2 mg/kg group. These results suggest that renal damage occurred in a dose-dependent manner only in mice treated with cisplatin. The liver tissues retained normal architecture in all groups, corresponding to no apparent tissue damage.

Together, these results show that dasatinib indeed exerts antitumor activity *in vivo* by inducing extensive intratumoral cell death, even in cases in which the tumor size without considering the inside contents appears to increase. Especially, dasatinib achieved tumor suppression that exceeded that by cisplatin under certain conditions, supporting its therapeutic potential.

### Tumor growth inhibition by the combination treatment of cisplatin and dasatinib *in vivo*

Cisplatin is indeed an effective drug in cancer chemotherapy. However, because of severe toxicities, limited patients are applicable for treatment by performance status^12^. Our data also show the cytotoxicity in kidney with clinically relevant dose (4 mg/kg, Fig. 4D). Therefore, the chemotherapy regimens which can keep the tumor suppressive effect and reduce the dose for cisplatin is critical issue clinically. As shown in Figure 4, dasatinib has effective for tumor suppression in OSCC likely without side effect because body weight reduction was not observed, raising the possibility that the decreased tumor suppressive effect through the reduced dose of cisplatin for decreasing the cytotoxicity can be compensated by combination treatment of dasatinib and cisplatin. Thus, we finally evaluated the possibility that co-administration of dasatinib and cisplatin can decrease the cisplatin dose and keep equivalent tumor suppressive effect without obvious side effect *in vivo*. The dosing schedule was identical to that used for the single-agent treatment (Fig. 4), and mice bearing subcutaneous tumors were assigned to three groups: control, dasatinib 30 mg/kg + cisplatin 4 mg/kg, and dasatinib 30 mg/kg + cisplatin 2 mg/kg (Fig. 5A).

**Figure 5.**
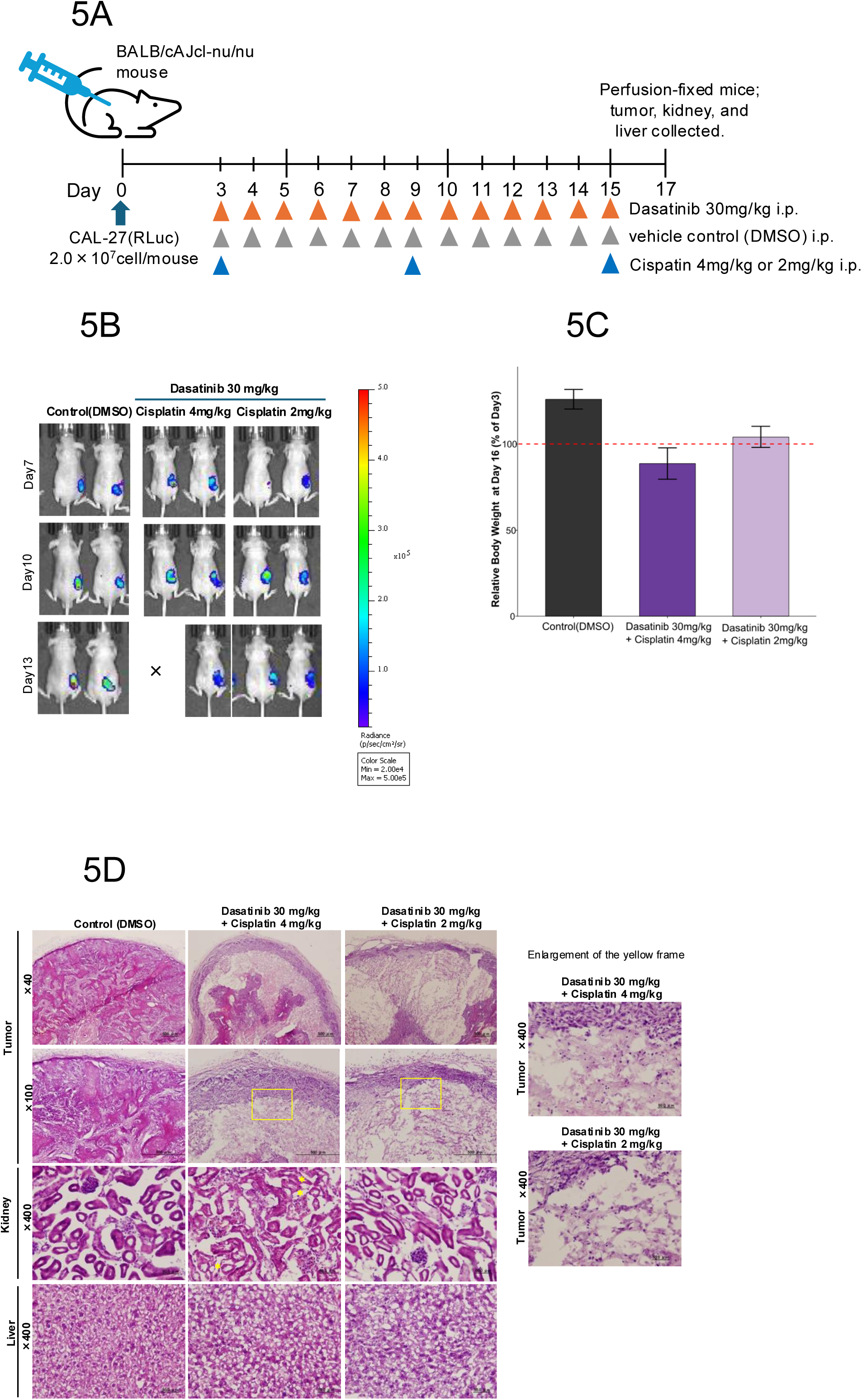
Tumor growth inhibition by combination treatment of cisplatin and dasatinib *in vivo*. **A** Experimental schedule for the combination treatment with dasatinib and cisplatin *in vivo*. CAL27-Rluc cells (clone 7) (2 × 10^7^ cells per mouse) were subcutaneously implanted into the right flank of 5-week-old male BALB/cAJcl nu/nu mice. Three days following implantation, the mice were randomly assigned to three treatment groups: vehicle control (DMSO), dasatinib (30 mg/kg) + cisplatin (4 mg/kg), and dasatinib (30 mg/kg) + cisplatin (2 mg/kg). (n=5–6). **B** IVIS bioluminescence live imaging of the tumor burden (additional mice are illustrated in Fig. S4). Bioluminescence signals were quantified 10 min following ViviRen injection according to the manufacturer’s instructions. For comparison, the same control image as in Figure S2 is shown again. **C** Body weight of nude mice normalized to Day 3 at the final Day 16 time point (mean ± s.d., n = 5–6). **D** Hematoxylin and eosin staining of the tumor, kidney, and liver sections. Mice were perfusion-fixed on day 17 and harvested tissues were processed to generate frozen sections for histological analysis. Representative images of each organ are indicated. Tumor histology: Control group showed solid squamous cell carcinoma. Dasatinib + cisplatin 4 mg/kg and dasatinib + cisplatin 2 mg/kg group exhibited a hollow, cyst-like structure with evidence of cell death within the lumen. Kidney histology: Control (DMSO) group displayed normal renal architecture. dasatinib + cisplatin 4 mg/kg group caused tubular epithelial degeneration and atrophy with focal rupture (*), while dasatinib + cisplatin 2 mg/kg groups induced mild tubular epithelial atrophy. Liver histology: All groups showed normal hepatic morphology without detectable abnormalities. Scale bars: 500 µm (×40, ×100); 50 µm (×400).

The data of tumor burden using IVIS bioluminescence live imaging showed substantially decreased tumor signals in both combination groups compared with those in the control group on Day13 (Fig. 5B, Fig. S4). Astonishingly, the antitumor effect was almost comparable between the high and low dose of cisplatin together with dasatinib. In contrast, body weight in the dasatinib 30 mg/kg + cisplatin 4 mg/kg group was intensely reduced compared to the control group, whereas the reduction of body weight was suppressed in the dasatinib 30 mg/kg + cisplatin 2 mg/kg group (Fig. 5C). These results suggested that reducing the cisplatin dose in combination treatment mitigate its toxicity with sustaining tumor growth inhibition.

Histological analysis of the excised tumors showed extensive intratumoral cell death in both combination groups, similar to that noted with dasatinib monotherapy, supporting an antitumor effect that was not reflected by tumor size alone (Fig. 5D). Regarding the cytotoxicity, liver sections revealed no apparent organ toxicity as similar as the results from the single-agent treatment, whereas kidney sections showed tubular epithelial degeneration and atrophy, as well as localized rupture in the dasatinib 30 mg/kg + cisplatin 4 mg/kg group, similar results to the monotherapy of cisplatin 4 mg/kg (Fig. 4D), but less in the dasatinib 30 mg/kg + cisplatin 2 mg/kg group. These results indicate that reducing the cisplatin dose and combining it with dasatinib diminished the nephrotoxicity associated with high-dose cisplatin, while maintaining its antitumor effect.

Collectively, these findings indicate that combination treatment with dasatinib and cisplatin improves the antitumor activity *in vivo*. Notably, reducing the cisplatin dose in combination with dasatinib treatment preserved the antitumor efficacy and may help attenuate treatment-related toxicity.

## Discussion

Herein, we examined the signaling pathway associated with tumorigenicity in OSCC (Fig. 1A), involvement of Src activity in broad oral cancer cell lines (Fig. 2B), sensitivity of OSCC cells to Src inhibitors (Fig. 2A and C, Table 1) and, molecular basis underlying this variability, and the therapeutic effect of combining Src inhibition with cisplatin in *in vitro* and *in vivo* models (Fig. 3 and 5). Particularly, dasatinib showed a distinctive antitumor effect, characterized by extensive intratumoral cell death, leading to a substantial decrease in the tumor burden *in vivo* (Fig. 4B and E). Additionally, the dasatinib and cisplatin combination markedly decreased tumor volume, even under cisplatin-dose-reduced conditions (Fig. 5), suggesting its potential to maintain antitumor efficacy while mitigating toxicity.

OSCC is a molecularly heterogeneous malignancy characterized by aberrant activation of multiple oncogenic signaling pathways, including EGFR, PI3K/AKT/mTOR, JAK/STAT, MET, Wnt/β catenin, and RAS/RAF/MAPK^34^. These pathways are closely linked to Src, which serves as a central signaling node that regulates diverse malignant phenotypes. Several FDA-approved Src inhibitors showed considerable variability in Src inhibitor sensitivity among the OSCC cell lines (Fig. 2A). This heterogeneity likely reflects the underlying molecular diversity of OSCC and may explain why therapies targeting a single pathway, including the Src pathway, often fail to achieve sufficient efficacy. Previous studies on HNSCC have attributed Src inhibitors’ limited efficacy to numerous factors *in vivo*, including the non-lethal nature of Src inhibition in some tumor cells, compensatory activation of alternative pathways such as PI3K/AKT and STAT3^22,26^, and differences in EMT status or cell adhesion molecules^27,28^. Consistent with these observations, our findings support the notion that the therapeutic response to Src inhibition in OSCC is strongly influenced by molecular diversity. Our data did not support a high phosphorylated-Src/total-Src ratio, although this has been proposed as a predictor of dasatinib sensitivity^30^. Although Src activation has been reported in 93.75% of OSCC tissues^36^, our results suggest that assessing the Src activation status and the degree of downstream pathway suppression are necessary for predicting Src inhibitor sensitivity. An especially striking finding in this study was that dasatinib induced extensive intratumoral cell death *in vivo*, resulting in a considerable decrease in viable tumor mass, even when the overall tumor size remained unchanged or increased (Fig. 4 and 5). IVIS data from *in vivo* imaging and histological analyses confirmed a clear dissociation between tumor size and tumor burden, showing an antitumor effect that could not be captured by size-based evaluation alone in some cases. From this perspective, dasatinib showed a possibility that tumor-suppressive activity is comparable to or exceeding that of cisplatin, indicating that the effect underscores its therapeutic potential at least in OSCC. Although dasatinib-induced intratumoral cavitation and cell death have not been previously reported, studies on bladder cancer cells have revealed that dasatinib suppresses the phosphorylation of the PI3K/Akt/mTOR pathway and activates apoptosis-and autophagy related proteins^37^. These mechanisms may contribute to cell death of CAL-27 noted in this study. Furthermore, inhibition of angiogenesis may also play a role in necrotic formation because Src activation is necessary for VEGF expression^38^. Whether these effects are specific to CAL-27 or represent a broader OSCC phenotype requires further *in vivo* investigations. Especially, this necrosis-inducing effect was almost not noted with bosutinib (Fig. S3E). Although this suggests a dasatinib-specific response, the current data are insufficient to determine its specificity, indicating that comparisons with additional Src inhibitors are required.

The dasatinib and cisplatin combination markedly decreased the tumor burden, even under cisplatin dose-reduced conditions, indicating its potential to maintain antitumor efficacy while decreasing toxicity (Fig. 5). This combination seemed to produce additive rather than synergistic effects *in vitro*, likely reflecting the independent actions of the inhibition of Src downstream signaling by dasatinib and the impairment of DNA repair mechanisms by cisplatin. Despite previous studies have reported that dasatinib suppresses cisplatin-induced Src activation^39^ and reverses Src-dependent cisplatin resistance^40^, such effects were not noted in the cisplatin-resistant OSCC cell line HO-1-u-1 (Fig. 3G). This may be owing to the Src-independent nature of cisplatin resistance in HO-1-u-1 and its low sensitivity to Src inhibitors. Nevertheless, our results suggest that combining Src inhibitors with cisplatin may enable dose reduction and mitigate toxicity (Fig 3 and 5), offering a clinically meaningful strategy for overcoming resistance and enhancing tolerability. Although bosutinib also showed antitumor activity *in vivo*, its pronounced toxicity, including loss of body weight, may limit its clinical applicability (Fig. S3). The considerable differences in toxicity profiles among the Src inhibitors underscore the criticality of careful drug selection when considering Src-targeted therapies for OSCC. Dasatinib showed a favorable balance between tumor reduction and tolerability, suggesting that it may be a more suitable Src inhibitor for clinical use.

This study has some limitations. First, Src inhibitors were ineffective in all the OSCC cell lines, and based on the seven cell lines examined, the molecular determinants of sensitivity could not be conclusively defined. Future studies incorporating a larger and more diverse panel, including patient-derived cell lines, are necessary for identifying the molecular features that predict the responsiveness of the Src inhibitor. Second, complete tumor disappearance could not be confirmed in any of the groups because the endpoint of this study was defined as visible tumor enlargement in mice. Importantly, dasatinib-treated tumors frequently showed extensive central cell death, raising the possibility that they may eventually collapse and regress, even though the time course of this process remains elusive. Further examination is necessary to optimize the combination strategies and treatment schedules. Nonetheless, FDA-approved Src inhibitors are orally administered and are generally less toxic than cytotoxic chemotherapies, leaving considerable room for their clinical application in settings such as neoadjuvant therapy or palliative treatment. In summary, this study reassessed the therapeutic potential of Src inhibition in OSCC and showed that Src inhibitors, especially dasatinib, are promising candidates, both as monotherapy and in combination with cisplatin. Dasatinib’s ability to decrease tumor burden and the potential for toxicity reduction through a reduction in cisplatin dose provide a foundation for developing new therapeutic strategies aimed at expanding treatment options and enhancing outcomes in patients with OSCC. Thus, our results provide new insights that may allow the establishment of future therapeutic strategies for OSCC.

## Methods

### Western blotting

Cells listed in Table S1 after each assay (Methods S1-S2) were lysed using M-PER Mammalian Protein Extraction Reagent with protease and phosphatase inhibitors (Table S2), mixed with an equal volume of 2× sodium dodecyl sulfate (SDS) buffer, and boiled at 90 °C for 10 min. Proteins were separated using SDS-polyacrylamide gel electrophoresis and transferred onto polyvinylidene fluoride membranes. Membranes were blocked and incubated with primary antibodies for 1 h at 25 ℃. After washing by Tris-buffered saline containing 0.05% Tween-20, the membranes were incubated with secondary antibodies for 1 h at 25 ℃ followed by protein bands detection using Clarity Western ECL Substrate and ChemiDoc Toch MP Imaging System (RRID:SCR_019037, Bio-Rad Laboratories).

### Cell viability assays

Cell viability in each assay (Method S3-S4) was evaluated using the CellTiter-Glo® 2.0 Reagent (#G9242, Promega, WI). Following drug treatment, CellTiter-Glo® 2.0 reagent was added at a 1:1 ratio to the culture medium, mixed for 2 min on an orbital shaker and incubated for 10 min at 25 ℃ to stabilize the luminescent signal. The luminescence was measured using a Tecan Spark microplate reader (RRID:SCR_021897, TECAN, Männedorf, Switzerland). All experiments were conducted in triplicates (n = 3).

### Animal experiments

CAL27-Rluc cells established by lentiviral vectors (Method S5) (clone 7; 2 × 10^7^ cells/mouse) were subcutaneously implanted into the right flank of 5-week-old male BALB/cAJcl nu/nu mice (RRID:IMSR_JCL:JCL:MID-0001, CLEA Japan, Tokyo, Japan). The mice were kept in a 12-hour light-dark cycle and were given free access to food and water. Three days following implantation, the mice were randomly assigned to each experimental group.

To detect Renilla luciferase activity corresponding to tumorigenicity (Method S6), ViviRen™ In Vivo Renilla Luciferase Substrate (#P1232, Promega) was dissolved in dimethyl sulfoxide at 10 mg/mL to prepare a stock solution, which was subsequently diluted 1,000-fold in PBS containing 0.1% bovine serum albumin. A total of 100 μL (1 mg) of the diluted substrate was intravenously administered. Mice were anesthetized with isoflurane, and bioluminescence imaging was conducted using an IVIS system (RRID:SCR_027425, Revvity Inc., MA). Bioluminescence signals were quantified 10 min following ViviRen injection according to the manufacturer’s instructions^41^. IVIS image analysis was performed on days 7 and 13 after subcutaneous tumor transplantation and on days 7, 10, and 13, respectively.

Tumor volumes were measured using calipers and calculated as L × W² × 0.52. Body weight was monitored every 2-3 d throughout the treatment period. At the end of the treatment, the mice were perfusion-fixed, and the tumor, kidney, and liver were harvested (N=5-6). All animal experiments were conducted in accordance with the Animal Experimentation Regulations of the Jichi Medical University. The Institutional Animal Care and Use Committee of Jichi Medical University reviewed and approved the study protocol. The study subsequently received final authorization from the University President.

### Statistical analysis

No statistical methods were used to predetermine sample size. Mouse studies were randomized; however, the investigators were not blinded to group allocation during the experiments or outcome assessment. Western blot analysis was independently performed three times. The 93 Src-associated chemical compounds screening and IC₅₀ analysis for the Src inhibitors were calculated from three independent CellTiter-Glo assays, and mean ± standard deviation (SD) was obtained using Microsoft Excel. For *in vitro* combination assays of the Src inhibitors and cisplatin, data are presented as mean ± SD from n = 3, and P values were calculated using paired t-tests in Microsoft Excel. Statistical significance was defined as follows: p < 0.01 **, p < 0.05 *, and p ≧ 0.05 ns. For *in vivo* experiments, tumor volume and body weight were analyzed using data from surviving mice at each time point (n = 5–6), and mean ± SD was calculated using R (version 4.5.2).

## Acknowledgements

We thank all of our laboratory members, especially Akemi Takada, who supported the handling of our budgets. This work was supported by AMED under Grant Number, JP23bm1323001, JMU Graduate Student Start-Up Award, and JST SPRING, Grant Number JPMJSP2183.

## Author Contributions

K.O. conceived and designed the study. Y.O. performed all the experiments and analyzed the data. Y.O. and K.O. interpreted experimental results. Y.O. and K.O. wrote the manuscript. Y.O., T.N., H.M., and K.O. edited the manuscript. All the authors have reviewed and approved the final manuscript.

## Data Availability Statement

The authors declare that the data supporting the findings of this study are available in the article and its Supplemental Information files. All relevant data are available from the authors upon reasonable request.

## References

1. Rumgay, H., Colombet, M., Ramos da Cunha, A., Filho, A.M., Warnakulasuriya, S., Conway, D.I., Chaturvedi, A., Virani, S., Lauby-Secretan, B., Carvalho, A.L., et al. (2026). Global incidence of lip, oral cavity, and pharyngeal cancers by subsite in 2022. CA Cancer J Clin 76.

2. Cancer Today https://gco.iarc.who.int/today/.

3. Chow, L.Q.M. (2020). Head and Neck Cancer. Reply. N Engl J Med 382, e57.

4. Liu, F., Chen, F., Huang, J., Yan, L., Wu, J., Qiu, Y., Zheng, X., Zhang, R., Lin, L., and He, B. (2017). Prospective study on factors affecting the prognosis of oral cancer in a Chinese population. Oncotarget 8, 4352–4359.

5. Kim, M.J., and Ahn, K.M. (2024). Prognostic factors of oral squamous cell carcinoma: the importance of recurrence and pTNM stage. Maxillofac Plast Reconstr Surg 46, 8.

6. Leemans, C.R., Braakhuis, B.J., and Brakenhoff, R.H. (2011). The molecular biology of head and neck cancer. Nat Rev Cancer 11, 9–22.

7. Mäkitie, A.A., Alabi, R.O., Pulkki-Råback, L., Almangush, A., Beitler, J.J., Saba, N.F., Strojan, P., Takes, R., Guntinas-Lichius, O., and Ferlito, A. (2024). Psychological Factors Related to Treatment Outcomes in Head and Neck Cancer. Adv Ther 41, 3489– 3519.

8. Blatt, S., Krüger, M., Sagheb, K., Barth, M., Kämmerer, P.W., and Al-Nawas, B. (2022). Tumor Recurrence and Follow-Up Intervals in Oral Squamous Cell Carcinoma. J Clin Med 11.

9. Ghanaati, S., Udeabor, S.E., Winter, A., Sader, R., and Heselich, A. (2025). Cancer Recurrence in Operated Primary Oral Squamous Cell Carcinoma Patients Seems to Be Independent of the Currently Available Postoperative Therapeutic Approach: A Retrospective Clinical Study. Curr Oncol 32.

10. Slieker, F.J.B., Rombout, D.A.A., de Bree, R., and Van Cann, E.M. (2022). Local recurrence and survival after treatment of oral squamous cell carcinoma of the maxilla: A systematic review and meta-analysis. Oral Surg Oral Med Oral Pathol Oral Radiol 133, 626–638.

11. Lefebvre, J.L. (2005). Current clinical outcomes demand new treatment options for SCCHN. Ann Oncol 16 Suppl 6, vi7–vi12.

12. Rabik, C.A., and Dolan, M.E. (2007). Molecular mechanisms of resistance and toxicity associated with platinating agents. Cancer Treat Rev 33, 9–23.

13. Atashi, F., Vahed, N., Emamverdizadeh, P., Fattahi, S., and Paya, L. (2021). Drug resistance against 5-fluorouracil and cisplatin in the treatment of head and neck squamous cell carcinoma: A systematic review. J Dent Res Dent Clin Dent Prospects 15, 219–225.

14. Lokich, J., and Anderson, N. (1998). Carboplatin versus cisplatin in solid tumors: an analysis of the literature. Ann Oncol 9, 13–21.

15. Gebre-Medhin, M., Adrian, G., Engström, P., Cange, H.H., Hammarstedt-Nordenvall, L., Reizenstein, J., Brun, E., Nyman, J., Abel, E., Friesland, S., et al. (2025). Chemoradiation therapy With Cisplatin Versus Cetuximab in Patients With Locoregionally Advanced Head and Neck Squamous Cell Cancer-Mature Results of the ARTSCAN III Trial. Int J Radiat Oncol Biol Phys 123, 681–690.

16. Ferris, R.L., Blumenschein, G., Fayette, J., Guigay, J., Colevas, A.D., Licitra, L., Harrington, K., Kasper, S., Vokes, E.E., Even, C., et al. (2016). Nivolumab for Recurrent Squamous-Cell Carcinoma of the Head and Neck. N Engl J Med 375, 1856– 1867.

17. Larkins, E., Blumenthal, G.M., Yuan, W., He, K., Sridhara, R., Subramaniam, S., Zhao, H., Liu, C., Yu, J., Goldberg, K.B., et al. (2017). FDA Approval Summary: Pembrolizumab for the Treatment of Recurrent or Metastatic Head and Neck Squamous Cell Carcinoma with Disease Progression on or After Platinum-Containing Chemotherapy. Oncologist 22, 873–878.

18. Irby, R.B., and Yeatman, T.J. (2000). Role of Src expression and activation in human cancer. Oncogene 19, 5636–5642.

19. Aleshin, A., and Finn, R.S. (2010). SRC: a century of science brought to the clinic. Neoplasia 12, 599–607.

20. Mandal, M., Myers, J.N., Lippman, S.M., Johnson, F.M., Williams, M.D., Rayala, S., Ohshiro, K., Rosenthal, D.I., Weber, R.S., Gallick, G.E., et al. (2008). Epithelial to mesenchymal transition in head and neck squamous carcinoma: association of Src activation with E-cadherin down-regulation, vimentin expression, and aggressive tumor features. Cancer 112, 2088–2100.

21. Cheng, S.J., Kok, S.H., Lee, J.J., Yen-Ping Kuo, M., Cheng, S.L., Huang, Y.L., Chen, H.M., Chang, H.H., and Chiang, C.P. (2012). Significant association of SRC protein expression with the progression, recurrence, and prognosis of oral squamous cell carcinoma in Taiwan. Head Neck 34, 1340–1345.

22. Yang, Z., Liao, J., Cullen, K.J., and Dan, H. (2020). Inhibition of IKKâ/NF-êB signaling pathway to improve Dasatinib efficacy in suppression of cisplatin-resistant head and neck squamous cell carcinoma. Cell Death Discov 6, 36.

23. Ammer, A.G., Kelley, L.C., Hayes, K.E., Evans, J.V., Lopez-Skinner, L.A., Martin, K.H., Frederick, B., Rothschild, B.L., Raben, D., Elvin, P., et al. (2009). Saracatinib Impairs Head and Neck Squamous Cell Carcinoma Invasion by Disrupting Invadopodia Function. J Cancer Sci Ther 1, 52–61.

24. Brooks, H.D., Glisson, B.S., Bekele, B.N., Johnson, F.M., Ginsberg, L.E., El-Naggar, A., Culotta, K.S., Takebe, N., Wright, J., Tran, H.T., et al. (2011). Phase 2 study of dasatinib in the treatment of head and neck squamous cell carcinoma. Cancer 117, 2112–2119.

25. Fury, M.G., Baxi, S., Shen, R., Kelly, K.W., Lipson, B.L., Carlson, D., Stambuk, H., Haque, S., and Pfister, D.G. (2011). Phase II study of saracatinib (AZD0530) for patients with recurrent or metastatic head and neck squamous cell carcinoma (HNSCC). Anticancer Res 31, 249–253.

26. Sen, B., Saigal, B., Parikh, N., Gallick, G., and Johnson, F.M. (2009). Sustained Src inhibition results in signal transducer and activator of transcription 3 (STAT3) activation and cancer cell survival via altered Janus-activated kinase-STAT3 binding. Cancer Res 69, 1958–1965.

27. Lee, S., Park, S., Ryu, J.S., Kang, J., Kim, I., Son, S., Lee, B.S., Kim, C.H., and Kim, Y.S. (2023). c-Src inhibitor PP2 inhibits head and neck cancer progression through regulation of the epithelial-mesenchymal transition. Exp Biol Med (Maywood) 248, 492–500.

28. Ortiz, M.A., Mikhailova, T., Li, X., Porter, B.A., Bah, A., and Kotula, L. (2021). Src family kinases, adaptor proteins and the actin cytoskeleton in epithelial-to-mesenchymal transition. Cell Commun Signal 19, 67.

29. Wu, Z., Doondeea, J.B., Gholami, A.M., Janning, M.C., Lemeer, S., Kramer, K., Eccles, S.A., Gollin, S.M., Grenman, R., Walch, A., et al. (2011). Quantitative chemical proteomics reveals new potential drug targets in head and neck cancer. Mol Cell Proteomics 10, M111.011635.

30. Levitt, J.M., Yamashita, H., Jian, W., Lerner, S.P., and Sonpavde, G. (2010). Dasatinib is preclinically active against Src-overexpressing human transitional cell carcinoma of the urothelium with activated Src signaling. Mol Cancer Ther 9, 1128–1135.

31. Wang, H., Lu, Y., Wang, M., Shen, A., Wu, Y., Xu, X., and Li, Y. (2022). Src inhibitor dasatinib sensitized gastric cancer cells to cisplatin. Med Oncol 40, 49.

32. Hanahan, D., and Weinberg, R.A. (2000). The hallmarks of cancer. Cell 100, 57–70.

33. Zhou, Y., Wang, L., Liu, M., Jiang, H., and Wu, Y. (2025). Oral squamous cell carcinoma: Insights into cellular heterogeneity, drug resistance, and evolutionary trajectories. Cell Biol Toxicol 41, 101.

34. Tan, Y., Wang, Z., Xu, M., Li, B., Huang, Z., Qin, S., Nice, E.C., Tang, J., and Huang, C. (2023). Oral squamous cell carcinomas: state of the field and emerging directions. Int J Oral Sci 15, 44.

35. Perše, M. (2021). Cisplatin Mouse Models: Treatment, Toxicity and Translatability. Biomedicines 9, 1406. 10.3390/biomedicines9101406.

36. Theocharis, S., Klijanienko, J., Giaginis, C., Alexandrou, P., Patsouris, E., and Sastre-Garau, X. (2012). FAK and Src expression in mobile tongue squamous cell carcinoma: associations with clinicopathological parameters and patients survival. J Cancer Res Clin Oncol 138, 1369–1377.

37. Ho, J.N., Byun, S.S., Kim, D., Ryu, H., and Lee, S. (2024). Dasatinib induces apoptosis and autophagy by suppressing the PI3K/Akt/mTOR pathway in bladder cancer cells. Investig Clin Urol 65, 593–602.

38. Gray, M.J., Zhang, J., Ellis, L.M., Semenza, G.L., Evans, D.B., Watowich, S.S., and Gallick, G.E. (2005). HIF-1alpha, STAT3, CBP/p300 and Ref-1/APE are components of a transcriptional complex that regulates Src-dependent hypoxia-induced expression of VEGF in pancreatic and prostate carcinomas. Oncogene 24, 3110–3120.

39. Ceppi, P., Papotti, M., Monica, V., Lo Iacono, M., Saviozzi, S., Pautasso, M., Novello, S., Mussino, S., Bracco, E., Volante, M., et al. (2009). Effects of Src kinase inhibition induced by dasatinib in non-small cell lung cancer cell lines treated with cisplatin. Mol Cancer Ther 8, 3066–3074.

40. Masumoto, N., Nakano, S., Fujishima, H., Kohno, K., and Niho, Y. (1999). v-src induces cisplatin resistance by increasing the repair of cisplatin-DNA interstrand cross-links in human gallbladder adenocarcinoma cells. Int J Cancer 80, 731–737.

41. Futami, M., Sato, K., Miyazaki, K., Suzuki, K., Nakamura, T., and Tojo, A. (2017). Efficacy and Safety of Doubly-Regulated Vaccinia Virus in a Mouse Xenograft Model of Multiple Myeloma. Mol Ther Oncolytics 6, 57–68.

