## Supplemental information for "Heterogeneous Sensitivity to Src Inhibitors in Oral Squamous Cell Carcinoma and Its Implications for Combination Therapy with Cisplatin"

### Supplementary information

#### Supplementary Methods:

Method S1: Cell culture
Method S2: Detection of phosphorylated proteins
Method S3: Drug dosage in cell viability assay
Method S4: $IC_{50}$ determination
Method S5: Lentiviral production and generation of Renilla luciferase (RLuc)-expressing CAL-27 cells
Method S6: Drug dosage in animal experiments
Method S7: Measurement of tumor volumes
Method S8: Hematoxylin and Eosin (H&E) staining

#### Supplementary Figures:

Figure S1. Drug Sensitivity of CAL27-Rluc Bulk Cells and Clones
Figure S2. Supplementary IVIS bioluminescence images corresponding to Fig. 4
Figure S3. The inhibition of tumor growth by single treatment of Dasatinib and Bosutinib in vivo
Figure S4. Supplementary IVIS bioluminescence images corresponding to Fig. 5

#### Supplementary Tables:

Table S1: Concentrations of Src inhibitors used for the Western blot analyses.
Table S2: Antibodies and reagents for Western blot.
Table S3: Src-associated chemical compounds and reagents used in this study.
Table S4: Concentrations of Src inhibitors used for the Western blot analyses.
Table S5: Concentrations of Src inhibitors used for the cell viability and combination treatment assays.

#### Supplementary Methods:

**Method S1: Cell culture**

Cells were maintained with indicated media (Supplementary Table S1) at 37°C in a humidified atmosphere containing 5% $CO_2$ and used within three months of the second passage.

**Method S2: Detection of phosphorylated proteins**

For phosphorylation analysis following Src inhibitor treatment, HSC-2, HSC-3, HSC-4, SAS, HO-1-u-1, CAL-27, and SCC-25 cells were treated with dasatinib, ponatinib, vandetanib, or bosutinib for 18 h. The cells were seeded in 12 well plates at densities that yielded approximately 80% confluence at the treatment time. Supplementary Table S4 provides the concentrations utilized for each cell line. Cells were harvested using the same lysis procedure described above, and protein samples were subjected to SDS-PAGE and western blotting, as previously described.

**Method S3: Drug dosage in cell viability assay**

Cells were seeded into Costar 96-well flat white plates (3917, Costar, NY) at the following densities: HSC-2 ($0.8 \times 10^4$), HSC-3 ($1.0 \times 10^4$), HSC-4 ($0.8 \times 10^4$), SAS ($2.0 \times 10^4$), HO-1-u-1 ($1.5 \times 10^4$), CAL-27 ($1.5 \times 10^4$), and SCC-25 ($0.7 \times 10^4$) cells per well, and allowed to adhere overnight. For the initial screen, CAL-27 cells were treated with a library of 93 Src-associated chemical compounds(10μM) for 36 h. HSC-2, HSC-3, HSC-4, SAS, HO-1-u-1, CAL-27, and SCC-25 cells were treated with the Src inhibitors dasatinib (100 nM), ponatinib (2 μM), vandetanib (10 μM), saracatinib (5 μM), PP2 (50 μM), bosutinib (5 μM), or cisplatin (10 μM) for 72 h. All compounds were diluted in a culture medium at the final concentration of DMSO with 1%, and cell viability was calculated relative to the 1% DMSO control.

**Method S4: $IC_{50}$ determination**

Cells were treated with serial dilutions of dasatinib, ponatinib, saracatinib, or bosutinib for 36 h. Cell viability was measured as described above. $IC_{50}$ values were calculated from dose-response curves using nonlinear regression in R (version 4.5.2). All experiments were conducted in triplicates (n = 3).

For combination treatment assays, cells were treated with cisplatin (10 μM or 5 μM), each Src inhibitor alone, and the combination of cisplatin (5μM) with each Src inhibitor for 36 h. Supplementary Table S5 provides the Src inhibitor concentrations used for each cell line.

Cell viability was calculated relative to the 1% DMSO control. Luminescence was measured as described above. All experiments were performed in triplicates (n = 3).

**Method S5: Lentiviral production and generation of Renilla luciferase (RLuc)-expressing CAL-27 cells**

RLuc-expressing CAL-27 (CAL27-Rluc) cells were generated by infecting parental CAL-27 cells with lentiviral particles produced by HEK293T cells. Lentiviruses were produced by co transfecting HEK293T cells with the packaging plasmids pCMV-VSVG -RSV-Rev (1 μg/μL, #RDB04393, RIKEN) and pCAG-HIVgp-RRE (1 μg/μL, #RDB04394, RIKEN), together with the transfer vector CSII-EGFP-P2A-RLuc-IRES-Bsd (Original: CSII-CMV-MCS-IRES2-Bsd, #RDB04385, RIKEN) at a ratio of 1:1:2. Viral infection of CAL-27 cells was conducted in the presence of polybrene at a 10 μg/mL final concentration. Seven independent CAL27-Rluc clones were established following infection and blasticidin selection. RLuc activity in each clone was confirmed using EnduRen live cell Renilla luciferase substrate (#E6482 Promega) according to the manufacturer's instructions. The drug responsiveness of the clones was assessed 36 h following treatment with Src inhibitors and cisplatin under the same conditions used for the Cell-Titer Glo® assay. Supplementary Fig. 1 shows the detailed screening results for the seven clones.

**Method S6: Drug dosage in animal experiments**

Six treatment groups based on their body weight (n = 5-6 in each group): vehicle control (DMSO), dasatinib alone (30 mg/kg), cisplatin alone (4 mg/kg), cisplatin alone (2 mg/kg), dasatinib (30 mg/kg) + cisplatin (4 mg/kg), and dasatinib (30 mg/kg) + cisplatin (2 mg/kg). Dasatinib and DMSO were dissolved in 10 mM Tris-HCl (pH 7.4), and cisplatin was dissolved in 0.9% NaCl. All drugs were intraperitoneally administered at a volume of 100 μL per mouse. Dasatinib and DMSO were administered once daily, whereas cisplatin was administered once weekly for two consecutive weeks. This dosing schedule was designed to approximate the clinical dosing frequencies in humans.

In a separate experiment conducted under identical implantation and monitoring conditions, the mice were assigned to three groups: vehicle control (DMSO), dasatinib alone (30 mg/kg), and bosutinib alone (50 mg/kg). Bosutinib was dissolved in 10 mM Tris HCl (pH 7.4) and intraperitoneally administered once daily at a volume of 100 μL per mouse, following the same dosing schedule as in the dasatinib experiment.

**Method S7: Measurement of tumor volumes**

Tumor volumes were measured using calipers and calculated as L × $W^2$ × 0.52. Body weight was monitored every 2-3 d throughout the treatment period. At the end of the treatment, the mice were perfusion-fixed, and the tumor, kidney, and liver were harvested (N = 5-6).

**Method S8: Hematoxylin and Eosin (H&E) staining**

Excised tumor, kidney, and liver tissues were fixed in 1% paraformaldehyde at 4 °C overnight. The tissues were subsequently cryoprotected by sequential incubation in 10% sucrose/PBS and 20% sucrose/PBS at 4 °C, each for 24 h. After cryoprotection, the tissues were embedded in OCT compound (Sakura Finetek, Tokyo, Japan) and snap-frozen in liquid nitrogen. Frozen blocks were sectioned at 8 μm thickness, and the sections were stained with H&E using standard procedures. The sections were post-fixed in 2% paraformaldehyde, stained with hematoxylin, rinsed under running tap water, counterstained with eosin, dehydrated using graded ethanol, cleared in xylene, and mounted with coverslips. Histological images were obtained using a light microscope.

### Figure S1

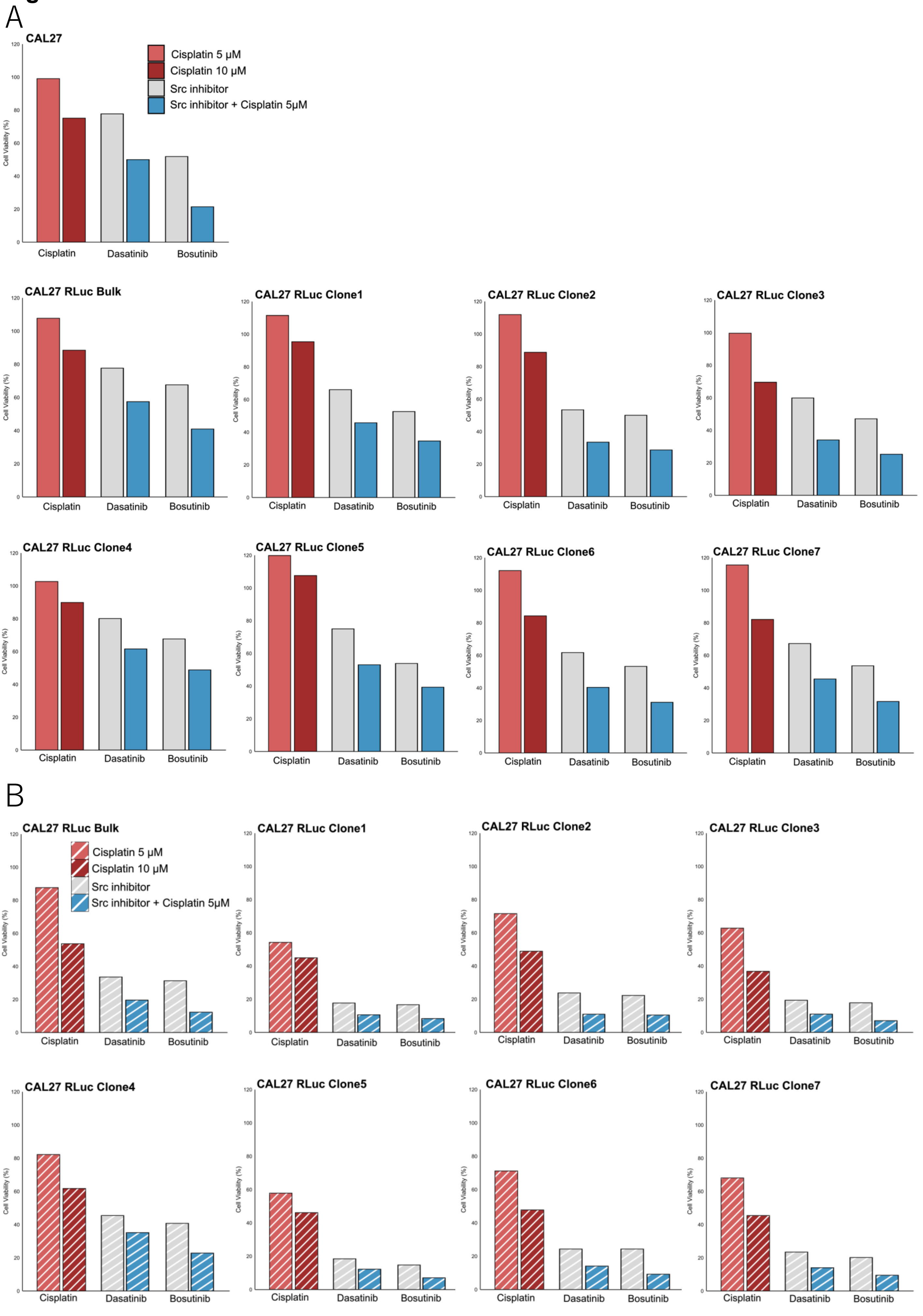
A
CAL27
Cisplatin 5 μM
Cisplatin 10 μM
Src inhibitor
Src inhibitor + Cisplatin 5μM
Cell Viability (%)
Cisplatin
Dasatinib
Bosutinib
CAL27 RLuc Bulk
CAL27 RLuc Clone1
CAL27 RLuc Clone2
CAL27 RLuc Clone3
CAL27 RLuc Clone4
CAL27 RLuc Clone5
CAL27 RLuc Clone6
CAL27 RLuc Clone7
B
CAL27 RLuc Bulk
Cisplatin 5 μM
Cisplatin 10 μM
Src inhibitor
Src inhibitor + Cisplatin 5μM
CAL27 RLuc Clone1
CAL27 RLuc Clone2
CAL27 RLuc Clone3
CAL27 RLuc Clone4
CAL27 RLuc Clone5
CAL27 RLuc Clone6
CAL27 RLuc Clone7

**Figure S1**
**Drug Sensitivity of CAL27-Rluc Bulk Cells and Clones.**

Drug sensitivity of stable Renilla luciferase (Rluc)-expressing CAL27 (CAL27-Rluc) bulk cells and clones 1–7 was evaluated 36 h following treatment using CellTiter-Glo (A) and EnduRen (B) assays.

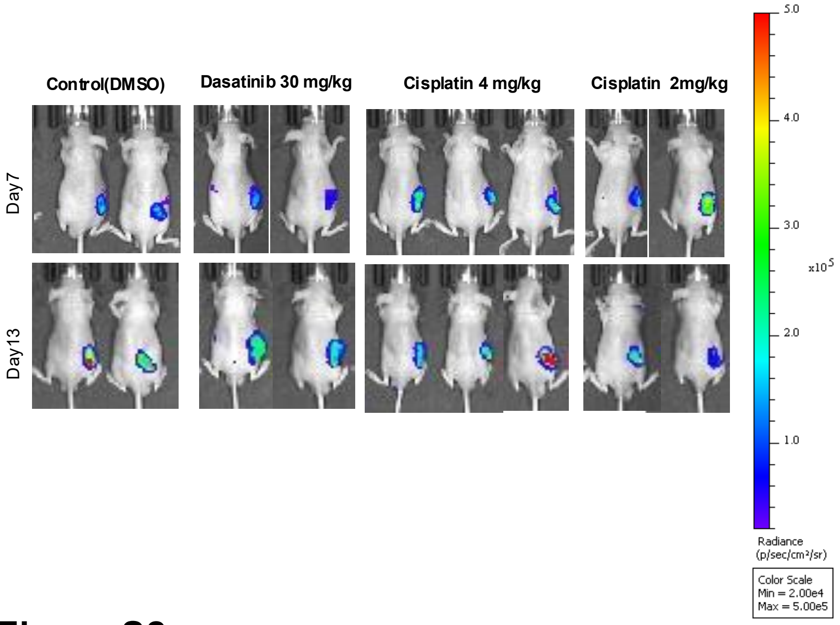


**Figure S2**
**Supplementary IVIS bioluminescence images corresponding to Fig. 4.**

Additional IVIS bioluminescence live images that are part of the total cohort shown in Fig. 4 (n = 5–6). Because Fig. 4B presents only three representative animals, the remaining images are provided here to document the full dataset used for analysis. All mice were imaged at the same time points as in Fig. 4, but these supplementary images were obtained from independent experimental runs.

**Figure S3**

**The inhibition of tumor growth by single treatment of Dasatinib and Bosutinib in vivo.**

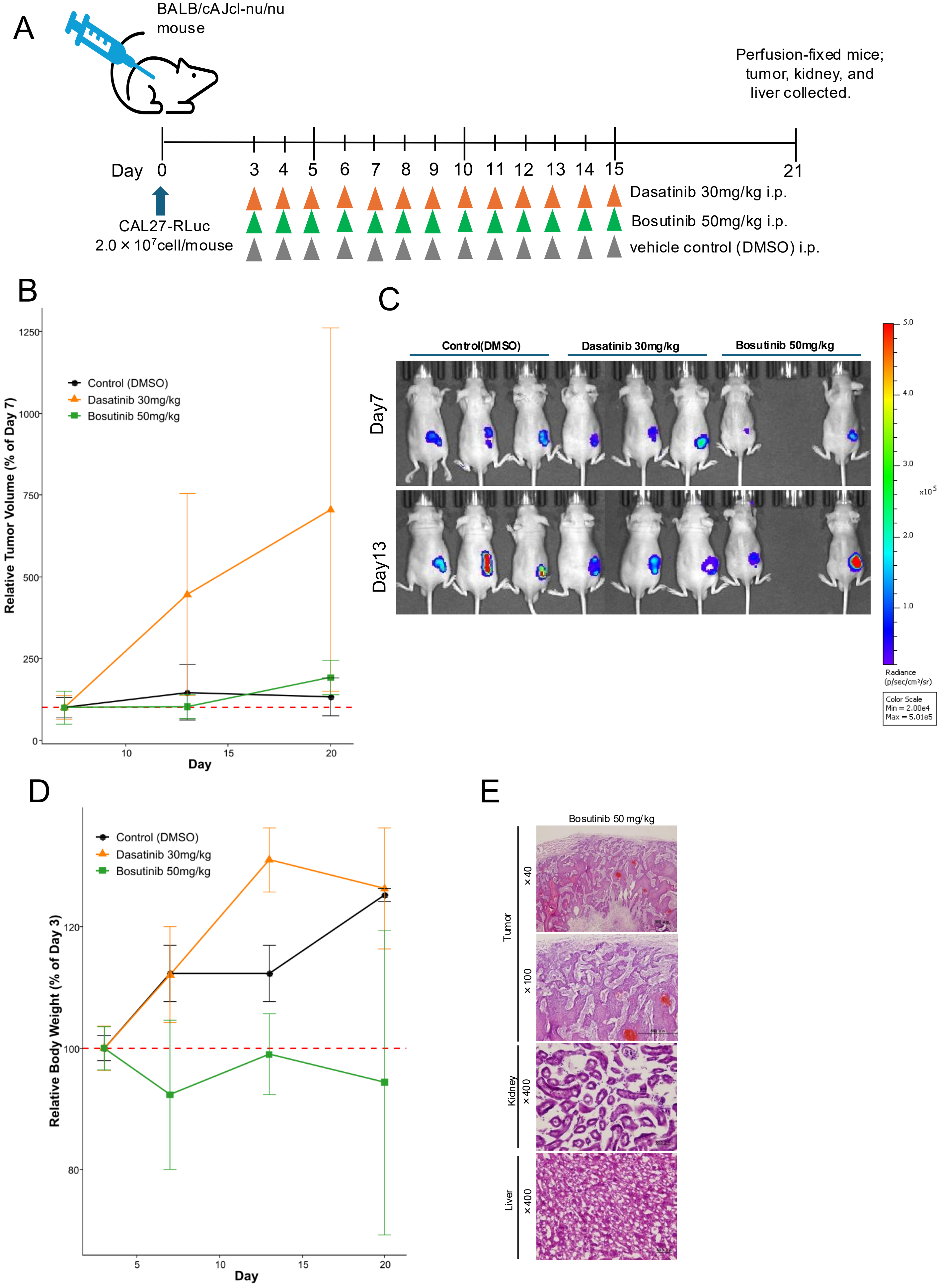

**Figure S3**
**The inhibition of tumor growth by single treatment of Dasatinib and Bosutinib in vivo.**

**A** Experimental schedule for single treatment with dasatinib and bosutinib in vivo. CAL27-Rluc cells (clone 7) (2 × 107 cells/mouse) were subcutaneously implanted into the right flank of 5-week-old male BALB/cAJcl nu/nu mice. Three days following implantation, the mice were randomly assigned to four treatment groups: vehicle control (DMSO), dasatinib alone (30 mg/kg), and bosutinib alone (50 mg/kg). Dasatinib, bosutinib, and DMSO were dissolved in 10 mM Tris-HCl (pH 7.4). All drugs were intraperitoneally administered at a volume of 100 µL per mouse. Dasatinib, bosutinib, and DMSO were administered once daily (n=3). **B** Tumor volumes normalized to Day 7 at the final Day 20 time point (mean ± standard deviation [s.d.], n = 2–3). Tumor volumes were measured using calipers and calculated as $L \times W^2 \times 0.52$. Measurements were conducted every 2–3 days. **C** IVIS bioluminescence live imaging of tumor burden. One of three mice in the bosutinib group died on day 6. **D** Body weights of nude mice from various treatment regimens implanted with CAL27-Rluc xenografts. Body weights are presented as mean ± standard deviation [s.d.] (n = 5–6) and were normalized to the wight on Day 3 for each mouse.(mean ± s.d., n = 2–3). **E** Hematoxylin and eosin staining of the tumor, kidney, and liver sections. Mice were perfusion-fixed on day 21 and the harvested tissues were processed to generate frozen sections for histological analysis. Representative images of each organ are presented. Tumor histology: Bosutinib group showed solid squamous cell carcinoma. Kidney histology: Bosutinib group displayed normal renal architecture. Liver histology: Bosutinib group showed normal hepatic morphology without detectable abnormalities. Scale bars: 500 µm (×40, ×100); 50 µm (×400).

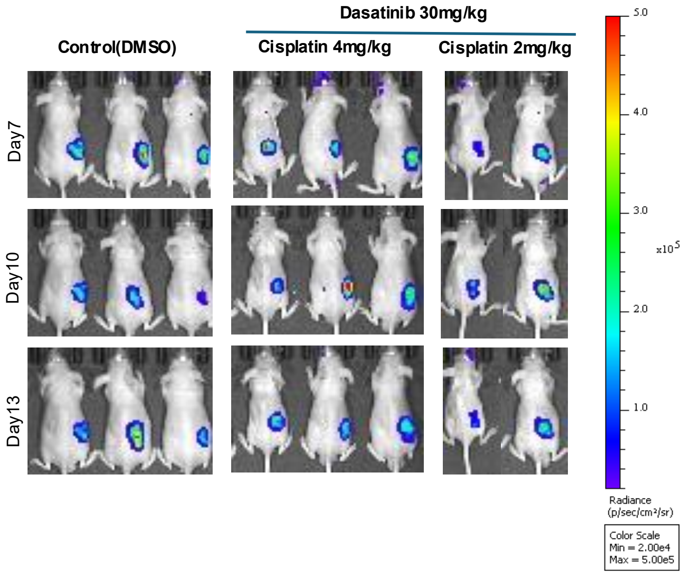


#### Figure S4
#### Supplementary IVIS bioluminescence images corresponding to Fig. 5.

Additional IVIS bioluminescence live images that are part of the total cohort shown in Fig. 5 (n = 5–6). Because Fig. 5B presents only two to three representative animals, the remaining images are provided here to document the full dataset used for analysis. All mice were imaged at the same time points as in Fig. 5, but these supplementary images were obtained from independent experimental runs. For comparison, the same control image as in Figure 4B is shown again.
